# Directed evolution of TurboID for efficient proximity labeling in living cells and organisms

**DOI:** 10.1101/196980

**Authors:** Tess C. Branon, Justin A. Bosch, Ariana D. Sanchez, Namrata D. Udeshi, Tanya Svinkina, Steven A. Carr, Jessica L. Feldman, Norbert Perrimon, Alice Y. Ting

## Abstract

Protein interaction networks and protein compartmentation underlie every signaling process and regulatory mechanism in cells. Recently, proximity labeling (PL) has emerged as a new approach to study the spatial and interaction characteristics of proteins in living cells. However, the two enzymes commonly used for PL come with tradeoffs – BioID is slow, requiring tagging times of 18-24 hours, while APEX peroxidase uses substrates that have limited cell permeability and high toxicity. To address these problems, we used yeast display-based directed evolution to engineer two mutants of biotin ligase, TurboID and miniTurbo, with much greater catalytic efficiency than BioID, and the ability to carry out PL in cells in much shorter time windows (as little as 10 minutes) with non-toxic and easily deliverable biotin. In addition to shortening PL time by 100-fold and increasing PL yield in cell culture, TurboID enabled biotin-based PL in new settings, including yeast, *Drosophila*, and *C. elegans*.

## Main text

Proximity labeling (PL) has emerged as an alternative to immunoprecipitation and biochemical fractionation for the proteomic analysis of macromolecular complexes, organelles, and protein interaction networks^1^. In PL, a promiscuous labeling enzyme is targeted by genetic fusion to a specific protein or subcellular region. Addition of a small molecule substrate, such as biotin, initiates covalent tagging of endogenous proteins within a few nanometers of the promiscuous enzyme (**Figure 1a**). Subsequently, the biotinylated proteins are harvested using streptavidin-coated beads and identified by mass spectrometry (MS).

**Figure 1.**
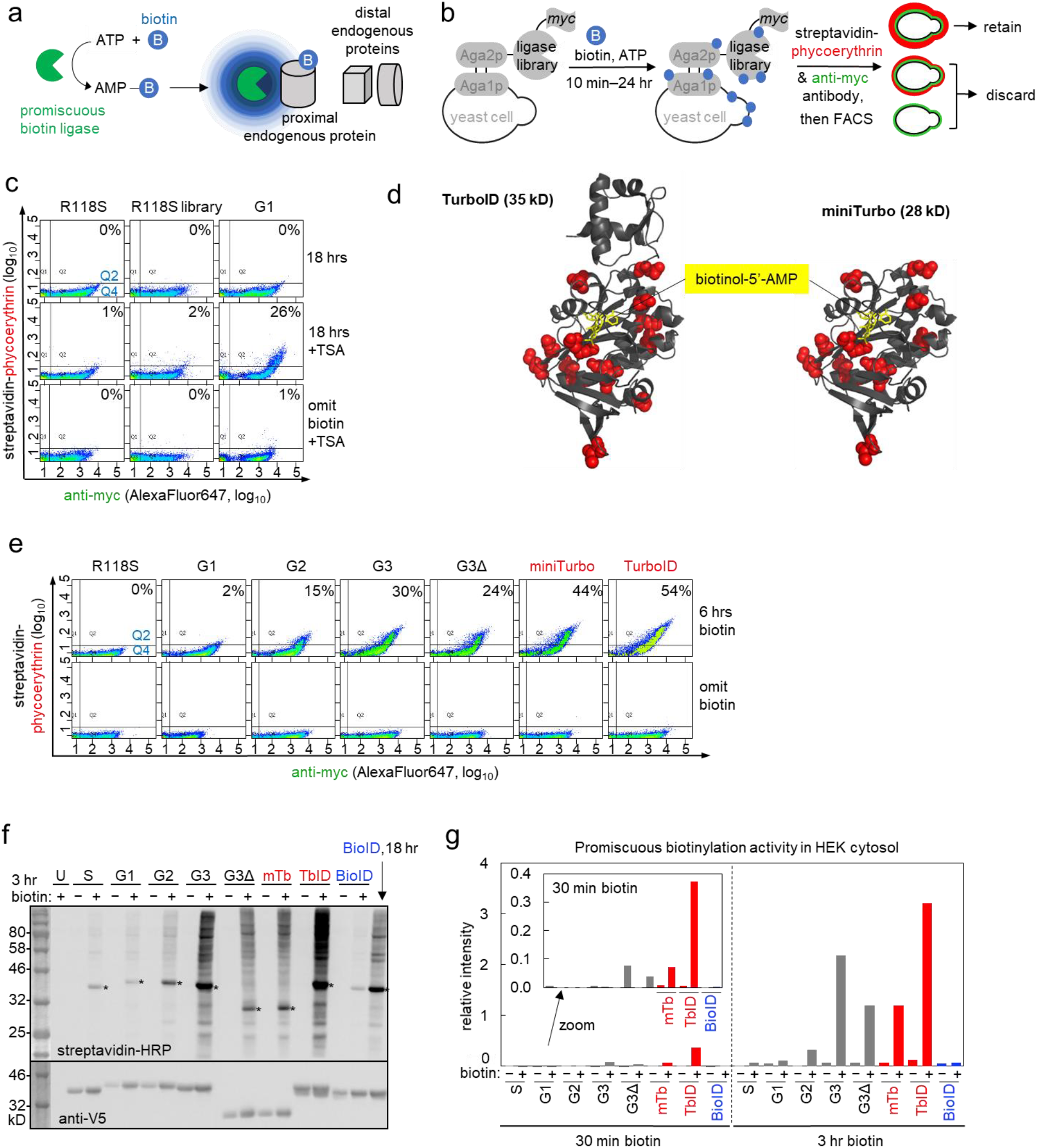
Directed evolution of TurboID. (**a**) Proximity-dependent biotinylation by promiscuous biotin ligases. Ligases catalyze the formation of biotin-5’-AMP anhydride, which diffuses out of the active site to biotinylate proximal endogenous proteins on nucleophilic residues such as lysine. (**b**) Yeast display-based selection scheme. A >10^7^ library of ligase variants is displayed on the yeast surface via fusion to mating protein Aga2p. All mutants have a C-terminal myc epitope tag. Biotin and ATP are added to the yeast library for between 10 minutes and 24 hours. Ligase-catalyzed promiscuous biotinylation is detected by staining with streptavidin-phycoerythrin and ligase expression is detected by staining with anti-myc antibody. Twodimensional FACS sorting enables enrichment of cells displaying a high ratio of streptavidin to myc staining. (**c**) Tyramide signal amplification (TSA) protocol improves biotin detection sensitivity on yeast. In the top row, the three indicated yeast samples were labeled with exogenous biotin for 18 hours then stained for FACS as in (b). The y-axis shows biotinylation extent, as measured by streptavidin-phycoerythrin staining intensity, and the x-axis quantifies ligase expression level. BirA-R118S is the template for the original ligase mutant library. G1 is the winning clone from the first generation of evolution. In the second row, after 18 hours of biotin treatment, the three yeast samples were stained with streptavidin-HRP, reacted with biotin-phenol^2^ to create additional biotinylation sites, then stained with streptavidin-phycoerythrin and anti-myc antibody before FACS. The third row is the same as the second row, but with the 18 hour biotin treatment omitted. Percentage of cells in upper right quadrant (Q2/(Q2+Q4)) indicated in top right of each graph. All plots display 10,000 cells.This experiment was performed once, but similar results under the same conditions have been repeated at least twice for each individual mutant in separate experiments. (**d**) Biotin ligase structure (PDB: 2EWN^26^) in gray with sites mutated in TurboID (left) and miniTurbo (right) colored red. The N-terminal domain (aa1-63) is also removed in the miniTurbo diagram. A nonhydrolyzable analog of biotin5’AMP, biotinol-5’-AMP, is shown in yellow stick. (**e**) FACS plots summarizing progress of directed evolution. Same presentation as in (c). G1G3 are the winning clones after generations 1-3 of directed evolution. G3Δ is G3 with the N-terminal domain deleted. miniTurbo (mTb) and TurboID (TbID) are our final promiscuous biotin ligases. All ligases were compared in parallel, with either 6 hours of 50 μM biotin + ATP incubation (top row), or growth in biotin deficient media (bottom row). All plots display 10,000 cells. This experiment was repeated one time (except for G3Δ and the “biotin omitted” conditions in “biotin depleted media’ which were performed only once under these conditions). (**f**) Comparison of ligase variants in HEK cytosol. The indicated ligases were expressed as NES (nuclear export signal) fusions in the HEK cytosol. 50 μM exogenous biotin was added for 3 hours, then whole cell lysates were analyzed by streptavidin blotting. Ligase expression detected by antiV-5 blotting. U, untransfected. Asterisks indicate ligase self-biotinylation. BioID labeling for 18 hours (50 μM biotin) shown for comparison (last lane). This experiment was performed once. (**g**) Quantitation of streptavidin blot data in (f) and from a 30 minute labeling experiment (blot shown in **Supplementary Figure 5a**). Quantitation is of promiscuous labeling bands and excludes self-biotinylation band. Sum intensity of each lane is normalized to that of BioID, 18 hours, which is set to 1.0.

Two enzymes are commonly used for PL: APEX2, an engineered variant of soybean ascorbate peroxidase^2,3^, and BirA-R118G (here, referred to as “BioID”), a point mutant of *E. coli* biotin ligase^4,5^. The main advantage of APEX2 is its speed: proximal proteins can be tagged in 1 minute or less, enabling dynamic analysis of protein interaction networks^6,7^. However, APEX labeling requires the use of H_2_O_2_, which is toxic to cells and difficult to deliver into live organisms without causing severe tissue damage. By contrast, BioID is attractive because of the simplicity of its labeling protocol and non-toxic labeling conditions – only biotin needs to be added to initiate tagging. These attributes have resulted in over 100 applications of BioID over the past 5 years, in cultured mammalian cells^5,8,9^, plant protoplasts^10^, parasites^11^–^19^, slime mold^20,21^, and mouse^22^. BioID has been used, for example, to map the protein composition of the centrosome-cilium interface^8^ and the inhibitory post-synaptic region^22^, each with nanometer spatial specificity.

The major disadvantage of BioID, however, is its slow kinetics, which necessitates labeling with biotin for 18-24 hours, and sometimes much longer^22^, to accumulate sufficient quantities of biotinylated material for proteomic analysis. This precludes the use of BioID for studying dynamic processes that occur on the timescale of minutes or even a few hours. Furthermore, low catalytic activity makes BioID difficult or impossible to apply in certain contexts - for example, in the ER lumen of mammalian cells, and in organisms such as yeast, worms, and flies.

A more efficient variant of BioID would greatly expand its utility, enabling the study of dynamic processes with minimal toxicity, and the extension of PL to new settings and organisms. Here, we report the directed evolution of *E. coli* biotin ligase to give two promiscuous variants, TurboID (35 kD) and miniTurbo (28 kD). Both are 7-26 fold more active than BioID, enabling proteomic labeling in just 10 minutes instead of the 18 hours commonly used for BioID. Furthermore, in 1 hour, TurboID can produce more biotinylated material than BioID produces in 18 hours. This enhanced activity also enabled us to perform biotin-based PL in new settings, including yeast, worm, and flies. Hence TurboID and miniTurbo broaden the scope of PL and should enable new discoveries related to spatial proteomes in living cells and organisms.

## Results

In an initial 2004 report, Cronan *et al.* tested three variants of *E. coli* biotin ligase and found that BirA-R118G (BioID) was the most promiscuous^4^. We started our engineering efforts by examining 7 other mutations at the R118 position, and found that R118S is ∽2-fold more active than BioID (**Supplementary Figure 1**). Hence, we selected BirA-R118S as our template for directed evolution.

In previous work, we used yeast display as our platform for directed evolution of APEX2^3^ and split-HRP^23^ enzymes. When coupled to Fluorescence Activated Cell Sorting (FACS), yeast display provides outstanding dynamic range, enabling separation of highly active enzyme mutants from moderately active ones. To employ this platform for directed evolution of promiscuous biotin ligase, we fused BirA-R118S or a library of R118S derived mutants (generated by error prone PCR) to the yeast cell surface mating protein Aga2p. We incubated the yeast cells with biotin and ATP for 18 hours to enable ligase-catalyzed PL to occur on the surface of each cell. We then stained the cells with streptavidin-phycoerythrin to visualize biotinylation sites, and anti-myc antibody to quantify ligase expression level, before two dimensional FACS sorting as shown in **Figure 1b**.

**Figure 1c** shows that biotinylation activity was undetectable by FACS for both BirA-R118S and our mutant library on the yeast surface, even with a labeling time of 18 hours. We could not proceed with a selection if none of the clones in our library showed activity above background. To increase the sensitivity of our assay, we explored amplification of biotin signal using the “tyramide signal amplification (TSA)” approach^24^. Instead of directly staining for biotinylation sites on the yeast surface with streptavidin-phycoerythrin, we first stained with streptavidin-horseradish peroxidase conjugate, then reacted with biotin-phenol to create more biotinylation sites^2^, and finally stained with streptavidinp-hycoerythrin (**Supplementary Figure 2a**). This approach made it possible to detect a small amount of signal over background (**Figure 1c**), and allowed us to begin directed evolution. We performed our first four rounds of selection using this amplification procedure, and reduced the labeling time gradually from 18 hours to 6 hours. After round four, activity of the pool was sufficient to omit the amplification step, and we again decreased our biotin labeling time from 3 hours to 1 hour (**Supplementary Figure 2b**).

Characterization of BirA mutant clones after six rounds of selection revealed that we had enriched some with high self-biotinylation activity. For example, clone R6-2 contains an E313K mutation that points directly into the BirA active site. We used mutagenesis to remove this lysine, and found that the resulting mutant, now called “G1”, still had ∽8-fold greater promiscuous biotinylation activity than our starting template, BirA-R118S (**Supplementary Figure 2b**). Hence we used G1, together with another clone, R6-1, as starting templates for a second library and a second generation of directed evolution. This time, to avoid enriching mutants with strong self-biotinylation, we treated the yeast, in some rounds of selection, with the reducing agent TCEP prior to streptavidin and myc staining, in order to cleave the Aga1p-Aga2p disulfide bonds that retain clones on the yeast surface (**Supplementary Figure 2c**). Six rounds of selection produced clone G2, with six mutations, as our best ligase mutant (**Supplementary Figure 2d**).

We continued with a third generation of directed evolution. Over four rounds, biotin labeling time was reduced to 10 minutes. The resulting winner, G3, was ∽17-fold more active than our starting template, BirA-R118S, but we noticed considerable activity on the yeast surface even without exogenous biotin addition (**Supplementary Figure 2e**). This suggests that G3 had evolved higher affinity for biotin and was able to utilize the low levels of biotin present in yeast culture media. An experiment in biotin-depleted media confirmed this hypothesis (**Supplementary Figure 2f**). We recognized the importance of remedying this problem because a BirA mutant that is capable of biotinylating prior to exogenous biotin addition prevents precise temporal control over the PL reaction. Starting from G3, we took two divergent paths.

For one path, we conducted a fourth generation of directed evolution, starting from a G3 mutant with its N-terminal domain (residues 1-63) deleted (“G3Δ”). **Supplementary Figure 2f** shows that this deletion alone reduces G3’s affinity for biotin, consistent with previous studies^25^. We conducted seven rounds of selection, including negative selections in which exogenous biotin was withheld (**Supplementary Fig. 2g**). The result was miniTurbo, with 12 mutations relative to wild-type BirA (**Supplementary Figure 2h** and **Figure 1d**).

In a second path, we performed a series of positive and negative selections on a library derived from full length G3 (**Supplementary Figure 2i**). One evolved mutation was retained, and we also carried over two mutations discovered in the evolution of miniTurbo. The result was TurboID, with 14 total mutations relative to wild-type BirA (**Figure 1d**).

On the yeast surface, we compared the activities of TurboID and miniTurbo to that of our starting template, BirA-R118S, and to various intermediate clones from our evolution (**Figure 1e** and **Supplementary Figure 3**). Each generation of evolution produced an increase in activity, while truncation of G3 to G3Δ decreased activity slightly. The most active mutants are miniTurbo and TurboID, with ∽17-fold and ∽19-fold higher activities than BirA-R118S on the yeast surface, respectively. Although TurboID has higher activity than miniTurbo, it also gives higher signal when exogenous biotin is omitted. Like G3, TurboID is able to utilize low levels of biotin present in regular yeast culture media.

We next tested whether these activity differences would replicate in a different context, the cytosol of cultured mammalian cells. We transfected NES (nuclear export signal)-tagged constructs into HEK293T cells, incubated with exogenous biotin for various lengths of time, terminated labeling by transferring cells to 4°C and washing away excess biotin (**Supplementary Figure 4**), and then lysed the cells and blotted the lysates with streptavidin-HRP. **Figures 1f, g** and **Supplementary Figure 5a** show that TurboID and miniTurbo are again the most active clones. For instance, TurboID gives more signal in 1 hour than the original BioID enzyme gives in 18 hours (**Figure 2a**). Both TurboID and miniTurbo produced easily detectable signal after just 10 minutes of incubation with exogenous biotin, while BioID signal was barely visible after 3 hours of biotin incubation (**Figure 1f** and **Supplementary Figures 5b-d**). Quantitation of Western blots, accounting for differences in ligase expression levels, showed that TurboID is 9-31 fold more active than BioID, while miniTurbo is 7-26-fold more active than BioID (**Figure 2b**). The decreased differences in activity between TurboID/miniTurbo and BioID at longer labeling times may result from saturation of available labeling sites.

**Figure 2.**
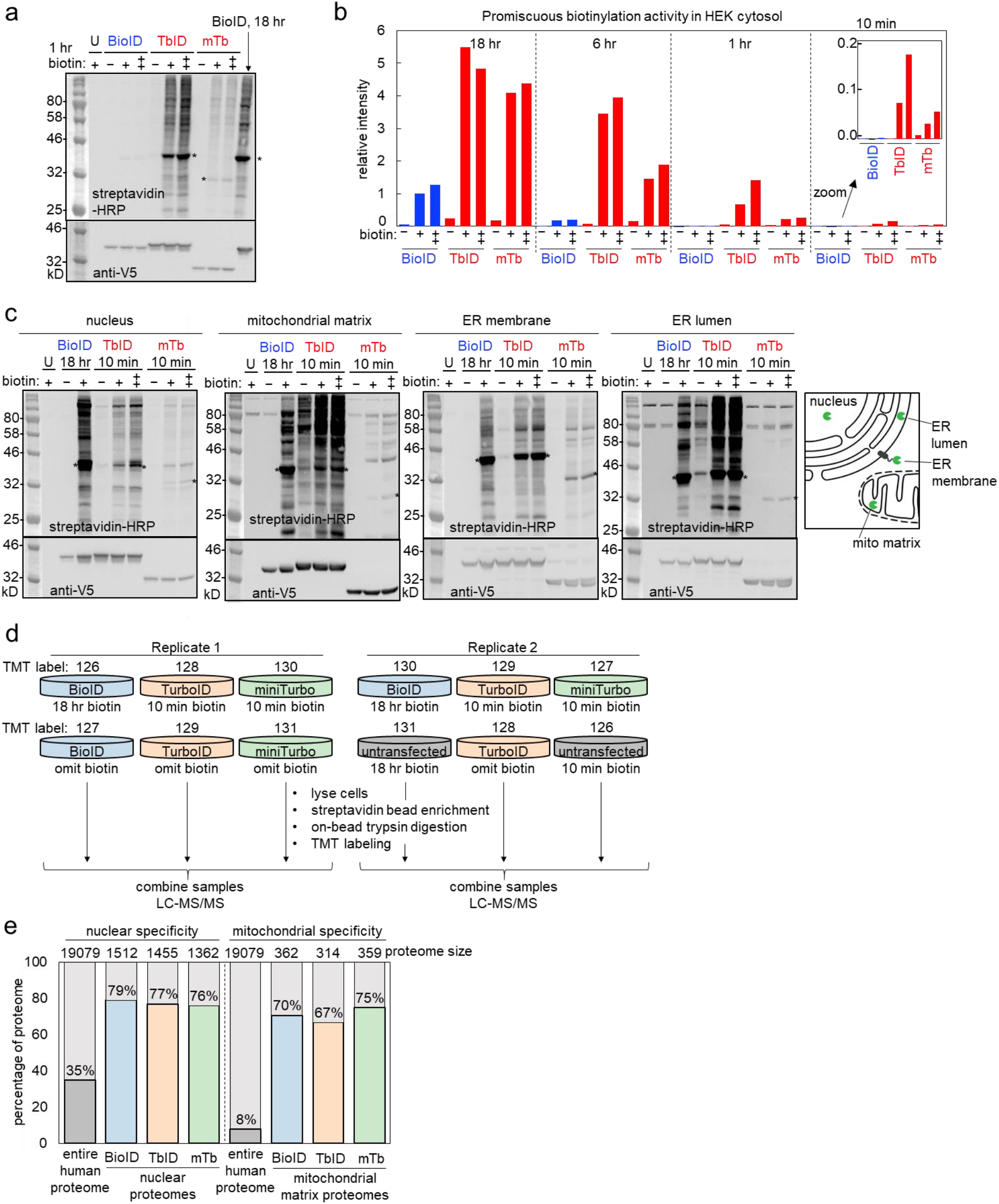
Characterization of TurboID and miniTurbo in mammalian cells. (**a**) Characterization of promiscuous biotinylation activity in HEK cells. Ligases were transiently expressed in the cytosol and 50 μM (+biotin) or 500 μM (++biotin) exogenous biotin was added for 1 hour. Whole cell lysates were analyzed by streptavidin blotting and ligase expression was detected by antiV-5 staining. U, untransfected. Asterisks indicate ligase selfbiotinylation. BioID 18 hour labeling is shown in rightmost lane for comparison. This experiment was performed once, but results under these conditions have been replicated for BioID three times. (**b**) Quantitation of streptavidin blot data in (a) in addition to multiple other experiments using 18 hr, 6 hr, and 10 min labeling times (blots shown in **Supplementary Figure 5bd**). Intensities are normalized to that of BioID, 18 hours, as in **Figure 1g**. (**c**) Comparison of promiscuous ligases in multiple HEK organelles. Each ligase was fused to a peptide targeting sequence (see Methods) directing them to the organelles indicated in the scheme at right. BioID samples were treated with 50 μM biotin for 18 hours. TurboID and miniTurbo samples were labeled for 10 minutes. Whole cell lysates were analyzed by streptavidin blotting. Ligase expression detected by antiV-5 blotting. + indicates treatment with 50 μM biotin; ++ indicates treatment with 500 μM biotin. U, untransfected. Asterisks indicate ligase selfbiotinylation. This experiment was repeated >3 times for nuclear constructs, twice for mitochondrial constructs, three times for ER membrane constructs, and once for ER lumen constructs. (**d**) Mass spectrometrybased proteomic experiment using TurboID, miniTurbo, and BioID. Experimental design and labeling conditions. HEK stably expressing TurboID or miniTurbo in the mitochondrial matrix (“mito”) or transiently in the nucleus (NLS) were treated with 500 μM exogenous biotin for 10 minutes. BioID samples expressed in the same manner were treated wth 50 μM biotin for 18 hours. After lysis, biotinylated proteins were enriched with streptavidin beads, digested to peptides, and conjugated to TMT labels. All six samples from each experiment were combined and analyzed by LCMS/MS. This experiment was performed once with two replicates for each construct. (**e**) Specificity analysis for each proteomic dataset. Graph indicates the fraction of each proteome with prior mitochondrial or nuclear annotation (according to GOCC^41,42^, MitoCarta^43^, or literature). Entire human proteome (according to GOCC^41,42^) shown for comparison. Number of proteins in each proteomic dataset shown across top.

Why do TurboID and miniTurbo have greater catalytic activity than BioID? Site-specific biotinylation catalyzed by wild-type BirA occurs via two half reactions: generation of the biotin-5’-AMP anhydride from biotin and ATP, followed by transfer onto a specific lysine of an acceptor peptide or protein^4^. It has been proposed that BirA-R118G (BioID) catalyzes promiscuous biotinylation by prematurely releasing biotin-5’-AMP into solution, for covalent tagging of nearby nucleophiles. **Figure 1d** shows that the 14 mutations in TurboID and miniTurbo are distributed throughout the ligase structure^26^, ^27^, with the majority far away (>10 Å) from the bound biotin-5’-AMP. Hence, if these mutations affect biotin-5’-AMP formation rate and/or affinity, it is likely via long-range and indirect mechanisms. Previous studies have shown that wild-type BirA dimerizes upon biotin-5’-AMP formation, and disruption of this dimer decreases affinity for biotin-5’-AMP^27^. Although every mutant in our study contains an A146 deletion that decreases dimerization^28^, it is possible that some of our evolved mutations further reduce dimerization. In addition, removal of the N-terminal domain is known to decrease biotin-5’-AMP affinity^25^. TurboID has two mutations at the junction between the N-terminal domain and the catalytic core, which may alter their structural relationship, leading to reduced biotin-5’-AMP affinity.

Different organelles have different pH, redox environments, and endogenous nucleophile concentrations, which may influence PL activity. We therefore compared TurboID, miniTurbo, and BioID in the nucleus, mitochondrial matrix, ER lumen, and ER membrane of HEK 293T cells (**Figure 2c**). While ligase activities varied by compartment, we were able to detect biotinylation by TurboID in 10 minutes, that was in some cases stronger than biotinylation by BioID in 18 hours. This was particularly striking in the ER lumen, where BioID activity was quite low, even after 18 hours, while TurboID gave robust labeling in 10 minutes. This activity difference is likely to be a consequence of our performing TurboID evolution in the oxidizing environment of the yeast secretory pathway.

Next, we tested TurboID and miniTurbo side-by-side with BioID in two quantitative proteomic experiments in the mammalian nucleus and mitochondrial matrix. For TurboID and miniTurbo, we supplied exogenous biotin for only 10 minutes; for BioID, we used 18 hours. Experiments were performed in replicate, alongside negative controls with ligases omitted or biotin omitted (**Figure 2d, Supplementary Figure 6**). After work up of the MS data as previously described^29^, we found that all three ligases gave proteomes of similar size and specificity (**Figure 2e**). The depth-of-coverage was somewhat smaller for TurboID and miniTurbo compared to BioID (**Supplementary Figure 6i**), but this could be a consequence of labeling for a 100-fold shorter time period.

Despite the widespread application of BioID, there has only been a single *in vivo* demonstration to date, in mice^22^, and certain cell types, such as yeast, are noticeably absent from published studies. We suspect that this is a consequence of BioID’s low catalytic activity, which makes it difficult or impossible to perform PL in certain cell types and organisms (for example, the mouse study required biotin addition for 7 days^22^). Though we carried out our directed evolution in the yeast secretory pathway, our starting template, BirA-R118S, gave almost undetectable signal in this context (**Figure 1c**). Consistent with this, we observed no promiscuous BioID activity at all in the yeast cytoplasm (**Figure 3a**). In contrast, TurboID and miniTurbo both gave robust labeling in this context. We also compared the three ligases in the bacterial cytosol. Here, TurboID and miniTurbo were also more active than BioID (**Figure 3b**).

**Figure 3.**
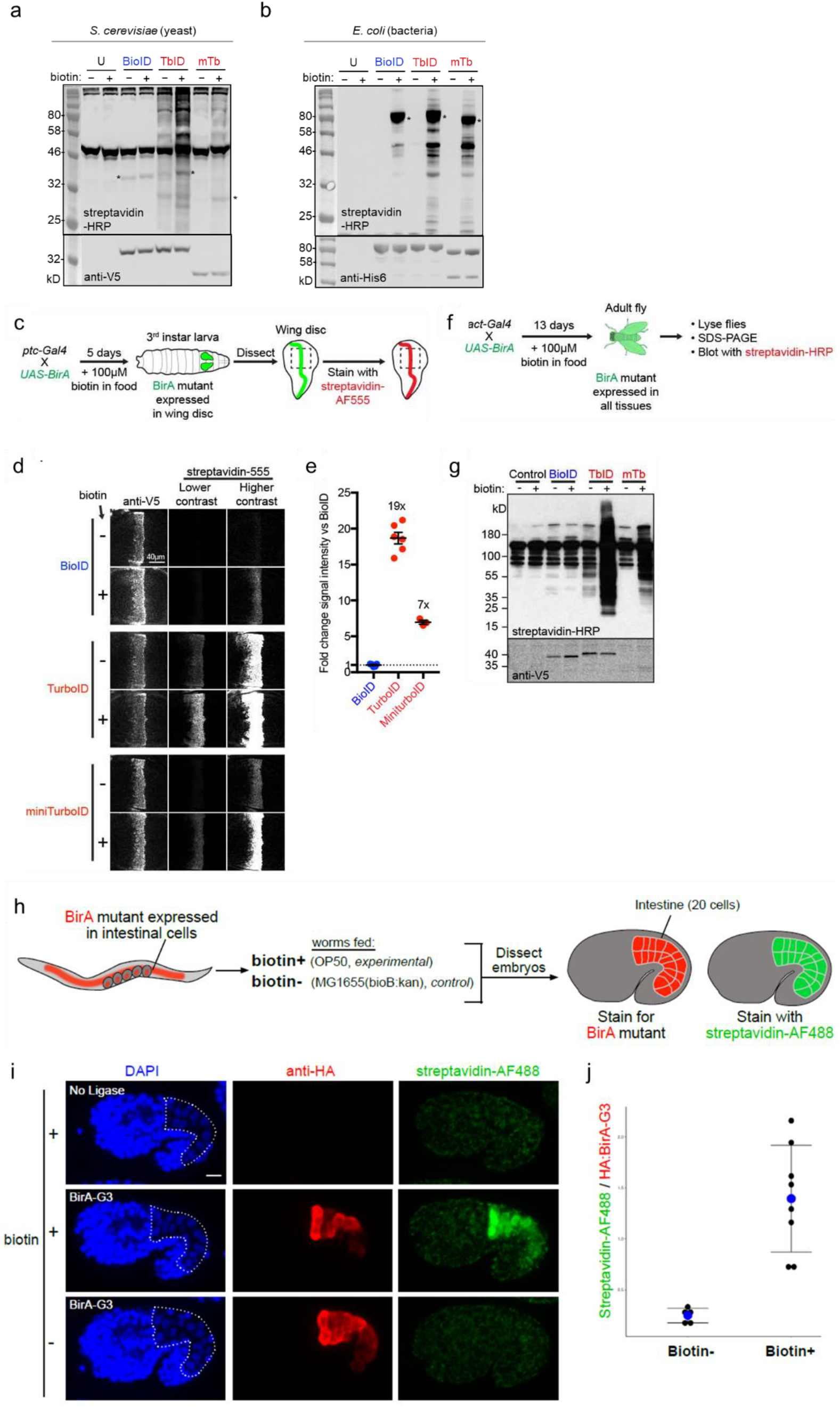
TurboID and miniTurbo in other species. (**a**) Yeast. Ligases were expressed in the cytosol of EBY100 *S. cerevisiae* and 50 μM exogenous biotin was added for 18 hours. Whole cell lysates were analyzed by streptavidin blotting. Ligase expression detected by antiV-5 blotting. U, untransfected. Asterisks indicate ligase self-biotinylation. This experiment was repeated once. (**b**) *E. coli.* Ligases fused at their N-terminal ends to maltose binding protein were expressed in the cytosol of BL21 *E. coli* and 50 μM exogenous biotin was added for 18 hours. Whole cell lysates were analyzed by streptavidin blotting. Ligase expression detected by antiHis6 blotting. U, untransfected. Asterisks indicate ligase self-biotinylation. This experiment was repeated once. (**c**) *D. melanogaster*. Schematic of tissuespecific labeling of endogenous proteins in fly wing disc. *Ptc-Gal4* induces expression of the indicated *UAS-BirA* enzyme in a strip of cells within the wing imaginal disc that borders the anterior/posterior compartments. Flies were fed with biotin-supplemented food for 5 days before dissecting and imaging. (**d**) Imaging of larval wing discs prepared as described in (c). Biotinylated proteins are detected by staining with streptavidin-AlexaFluor555, and ligase expression is detected by antiV-5 staining. Panels show the pouch region of the wing disc, indicated by the dashed line in (c). Scale bar, 40 μm. Each experimental condition has at least three technical replicates, one representative image is shown. This experiment was performed once. Similar experiments using temporally controlled *ptc-Gal4* expression, or an earlier version of TurboID (G3), showed similar results (not shown). (**e**) Quantitation of signal intensity of streptavidin-AlexaFluor555 in (d). Error bars in s.e.m. Average foldchange shown as text above plot lane. Sample size values (n) from left column to right: 5, 6, 3. (**f**) *D. melanogaster*. Scheme of ubiquitous labeling of endogenous proteins in flies. *Act-Gal4* drives expression of the indicated *UAS-BirA* enzyme in all cells at all developmental time points. Flies were provided food supplemented with 100 μM biotin for 13 days after egg deposition, then the whole body of adult flies were lysed. (**g**) Western blotting of fly lysates prepared as described in (e). Biotinylated proteins detected by blotting with streptavidin-HRP, ligase expression detected by antiV-5 blotting. In control sample, *Act-Gal4* drives expression of *UAS-luciferase* in all cells at all developmental time points. This experiment was repeated once. (**h**) *C. elegans.* Scheme of tissuespecific labeling of endogenous proteins in the *C. elegans* embryonic intestine. *ges1p* promoter drives expression of BirA mutant G3 in the intestine beginning approximately 150 minutes after the first cell cleavage. Transgenic strains are fed *OP50* bacteria (biotin+), or biotin-auxotrophic bacteria (*MG1655bioB:kan*; biotin-) for two generations to deplete excess biotin. Embryos are then assayed 6-7 hours into development, making the biotin labeling window approximately 4 hours. (**i**) Imaging of *C. elegans* embryos prepared as described in (h). Biotinylated proteins are detected by staining with streptavidin-AF488, and BirAG3 expression is detected by anti-HA staining. Intestine is outlined by a white dotted line. Scale bars, 5 μm. This experiment was performed once with at least 5 biological replicates. (**j**) Quantitation of mean ratios of streptavidin pixel intensity to antiHA intensity for biotin+ (n = 8, mean = 1.3939 ± 0.5236), or biotin - (n = 5, mean = 0.2469 ± 0.0726) embryos. Wilcoxon rank sum test was used to determine the difference between the two populations, *p* = 0.007937. Mean is shown as a large blue dot for each condition.

Two model organisms, *Drosophila* and *C. elegans*, are frequently used for biological studies due to their genetic tractability. In principle, they are also well-suited for BioID because biotin can be easily delivered to various organ systems through food^30^. Yet no published studies document the use of BioID in either organism. Here, we sought to test the applicability of biotin-based PL to these animals. In *Drosophila*, we expressed BioID, TurboID, or miniTurbo either ubiquitously, or selectively in the larval wing disc, which gives rise to the adult wing. After 5-13 days of feeding on biotin-containing food, we stained dissected wing discs with streptavidin-fluorophore (**Figures 3c-e**), or lysed the adult flies and ran a streptavidin blot (**Figures 3f, g**). In **Figures 3c-e**, TurboID and miniTurbo signals are 19-fold and 7-fold higher, respectively, than BioID signal in the wing disc. Consistent with our observations in HEK 293T cells, TurboID also gives some low signal in flies fed regular food (without biotin supplementation).

In the fly streptavidin blots, BioID activity is undetectable, whereas TurboID and miniTurbo both give robust biotinylation signal (**Figure 3f, g**). The absence of detectable BioID signal here versus when the ligase is expressed in a specific tissue may be due to endogenous biotinylated proteins drowning out specific signal in the streptavidin blot.

Separately, we performed adult survival and wing morphology assays to check for possible toxic effects of BioID, TurboID, or miniTurbo expression in flies (**Supplementary Figure 7**). When expressed ubiquitously, BioID and miniTurbo were non-toxic, but TurboID flies showed decreased survival and were smaller in size when grown without biotin supplementation. We hypothesize that TurboID consumes all the biotin, effectively biotin starving cells, when expressed ubiquitously. In support of this, toxicity can be rescued by supplementing the fly’s food with exogenous biotin. Furthermore, none of the ligases were toxic when expressed in the wing disc, since adult wings were observed to be normal in morphology and size.

In *C. elegans*, we tested our TurboID precursor, BirA-G3, in the embryonic intestine, a simple epithelial tube composed of 20 cells. The intestinal lineage is specified 35 minutes after the 2-cell stage of the embryo and intestinal cells begin to differentiate about 4 hours later^31,32^. Thus, performing PL during the early stages of intestinal development requires a ligase with sufficient activity to label within a window of only a few hours. We expressed BirA-G3 early in the intestinal lineage (approximately 150 min after the first cleavage) and assayed biotinylation activity approximately 4 hours later. BirA-G3 showed robust biotinylation activity in embryos from worms fed biotin-producing bacteria (**Figures 3h-j**). In contrast, embryos from worms fed bacteria unable to biosynthesize biotin showed the same low level of biotinylation in the intestinal cells as embryos that lacked the G3 ligase (**Figures 3h-j**).

## Discussion

In summary, we have used yeast display-based directed evolution to engineer two BirA variants, TurboID and miniTurbo, with much greater promiscuous biotinylation activity than the original BioID enzyme, BirA-R118G. BioID has already had tremendous impact in the proteomics field, enabling sub-compartment mapping^8,22^ and protein-protein interaction^19,33^ discovery with greater specificity and sensitivity than traditional approaches, such as biochemical fractionation and immunoprecipitation, allow. The introduction of these two new enzymes, which enable live cell proximity biotinylation with greater signal in shorter time windows – as little as 10 minutes instead of the 18-24 hours typically used for BioID – should further expand the scope of this important methodology.

We engineered two enzymes in this study instead of one, because they each have unique properties and tradeoffs. TurboID is the most active, and should be used when the priority is to maximize biotinylation yield and sensitivity/recovery. However, in many contexts, we observe a small amount of promiscuous biotinylation before exogenous biotin is supplied, indicating that TurboID can utilize the low levels of biotin present in cells and organisms grown in typical biotin-containing media/food. Nearly all eukaryotes import biotin, as they cannot biosynthesize their own^34^. Interestingly, bacteria, which can make their own biotin, did not give TurboID background before exogenous biotin addition (**Figure 3b**). Perhaps bacteria have lower levels of free biotin due to feedback regulation of its synthesis.

If, on the other hand, the priority is to precisely restrict promiscuous biotinylation to a specific window of time, then miniTurbo is recommended over TurboID. miniTurbo is not as active as TurboID, but it gives much less background than TurboID in the absence of exogenous biotin addition. Another benefit of miniTurbo is that it is 20% smaller than TurboID (28 versus 35 kD), which may decrease the probability of negative impact on the trafficking and/or function of the proteins to which it is fused.

In addition to decreasing the time window of labeling and increasing signal, TurboID and miniTurbo enable BirA-based proximity labeling in new contexts that we showed are problematic for the original BioID enzyme. We believe that the beneficial properties of TurboID and miniTurbo arise from the fact that they were evolved in the yeast secretory pathway at 30°C (the normal culturing temperature for yeast), while wild-type BirA normally functions in the cytosol of *E. coli* at 37°C^4^. Hence, TurboID was much more active than BioID in the mammalian ER lumen, and TurboID gave robust biotinylation in the yeast cytosol (at 30°C), where BioID activity was undetectable. Our BirA variants were also efficacious in flies which grow at 25°C and in worms which grow at 20°C.

Despite the popularity of enzyme-catalyzed proximity labeling, there have been very few *in vivo* applications to date. BioID has only been used in the mouse brain for mapping of the inhibitory post-synapse, where biotin was supplied by IP injection for 7 days^22^, and in a xenograft model, where biotin was supplied by IP injection for 2 days^33^. APEX peroxidase has been used in three in vivo studies, but in each case, genetic modification to compromise cuticle integrity^35,36^ or manual dissection of tissue had to be performed^37^ to deliver APEX chemical substrates to the relevant cells. APEX also relies on H_2_O_2_ which is toxic. Hence TurboID and miniTurbo expand the possibilities for non-toxic but rapid proximity biotinylation, with facile substrate delivery, in in vivo systems.

Our lab has previously used yeast display-based directed evolution to improve the catalytic efficiency of APEX2 and split HRP enzymes. However, here we faced new challenges that required a number of innovations. First, starting signal was far too low, and required the development of a signal amplification procedure. TSA has been used on fixed mammalian cells for fluorescence microscopy applications^38^, but not, as far as we are aware, on live yeast cells, or for FACS. Second, to distinguish promiscuous biotinylation activity from self-biotinylation activity, we developed a strategy to remove ligases from the yeast surface prior to staining and FACS. Third, we found it essential to implement negative selections to eliminate ligases with increased biotin affinity that would enable them to use the low levels of biotin present in normal media. These strategies may be beneficial to others seeking to use the yeast display platform to evolve new enzymatic activities.

Recently, a BioID variant from *Aquifex aeolicus* was reported, called BioID2^39^. BioID2 is 25% smaller than BioID, and more active at higher temperatures, but not claimed (or shown) to be faster or more catalytically efficient than BioID. One follow-up study used BioID2 for proteomic mapping of the inner nuclear membrane and employed a biotin tagging time of 16 hours^40^. BioID2 also has higher biotin affinity than BioID, described by the authors as an advantage, but this results in biotinylation activity in the absence of exogenous biotin addition, which prevents precise temporal control over labeling.

## Acknowledgements

FACS experiments were performed at the Koch Institute Flow Cytometry Core (MIT) and Stanford Shared FACS Facility. S. Han (Stanford) synthesized neutravidin-AlexaFluor647. We thank I. Droujinine (Harvard) for advice on biotin labeling in *D. melanogaster.* This work was supported by NIH R01-CA186568 (to A. Y. T.), Howard Hughes Medical Institute HCIA grant (to N.P.), NIH New Innovator Award DP2GM119136-01 (to J. L. F.) and Howard Hughes Medical Institute Collaborative Innovative Award (to S.A.C. and A.Y.T.). A.Y.T. is a Chan Zuckerberg Biohub investigator. T.C.B. was supported by Dow Graduate Research and Lester Wolfe Fellowships. J.A.B. was supported by Damon Runyon Post-Doctoral Fellowship. A.D.S. was supported by NIH Training Grant 2T32GM007276.

## Author contributions

T.C.B. and A.Y.T. designed the research and analyzed all the data except those noted. T.C.B. performed all experiments except those noted. T.C.B., A.Y.T., N.D.U. and S.A.C. designed the proteomics experiments. T.C.B. prepared the proteomic samples. N.D.U. and T.S. processed the proteomic samples and performed mass spectrometry. J.A.B. performed *D. melanogaster* experiments. J.A.B. and N.P. analyzed *D. melanogaster* data. T.C.B., A.Y.T., A.D.S., and J.L.F. designed the *C. elegans* experiments. A.D.S. performed *C. elegans* experiments. A.D.S. and J.L.F. analyzed *C. elegans* data.

## Supporting Figures

**Supplementary Figure 1.**
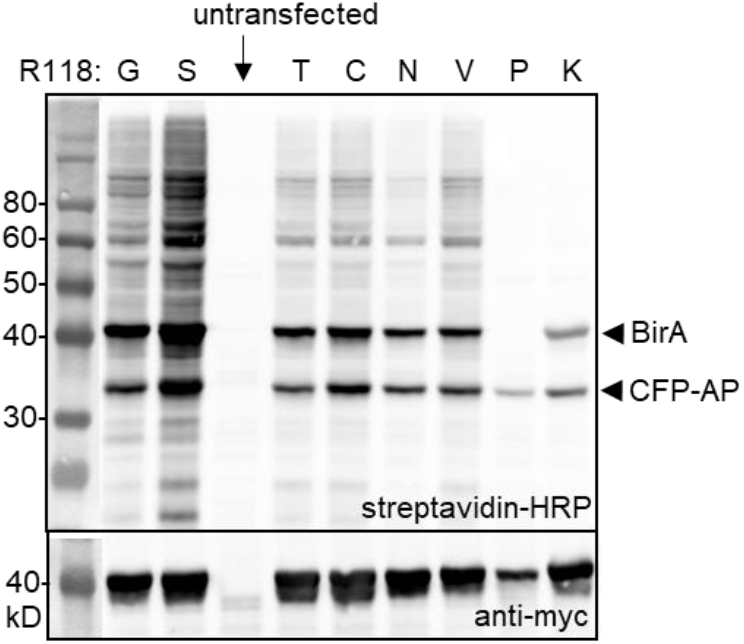
Testing mutations at R118 of BirA. BirA ligases with the indicated mutations were transiently expressed as NES (nuclear export signal) fusions in the HEK cytosol. All samples were cotransfected with AP-CFP (acceptor peptide fused to cyan fluorescent protein), which is site-specifically biotinylated by BirA^44^. 50 μM exogenous biotin was added to the cells for 18 hours, then whole cell lysates were analyzed by streptavidin-HRP blotting. Ligase expression was detected by anti-myc blotting. Self-biotinylation bands are indicated (“BirA”). This experiment was performed once, but comparison of BirA-R118G to BirA-R118S under these conditions was repeated once.

**Supplementary Figure 2.**
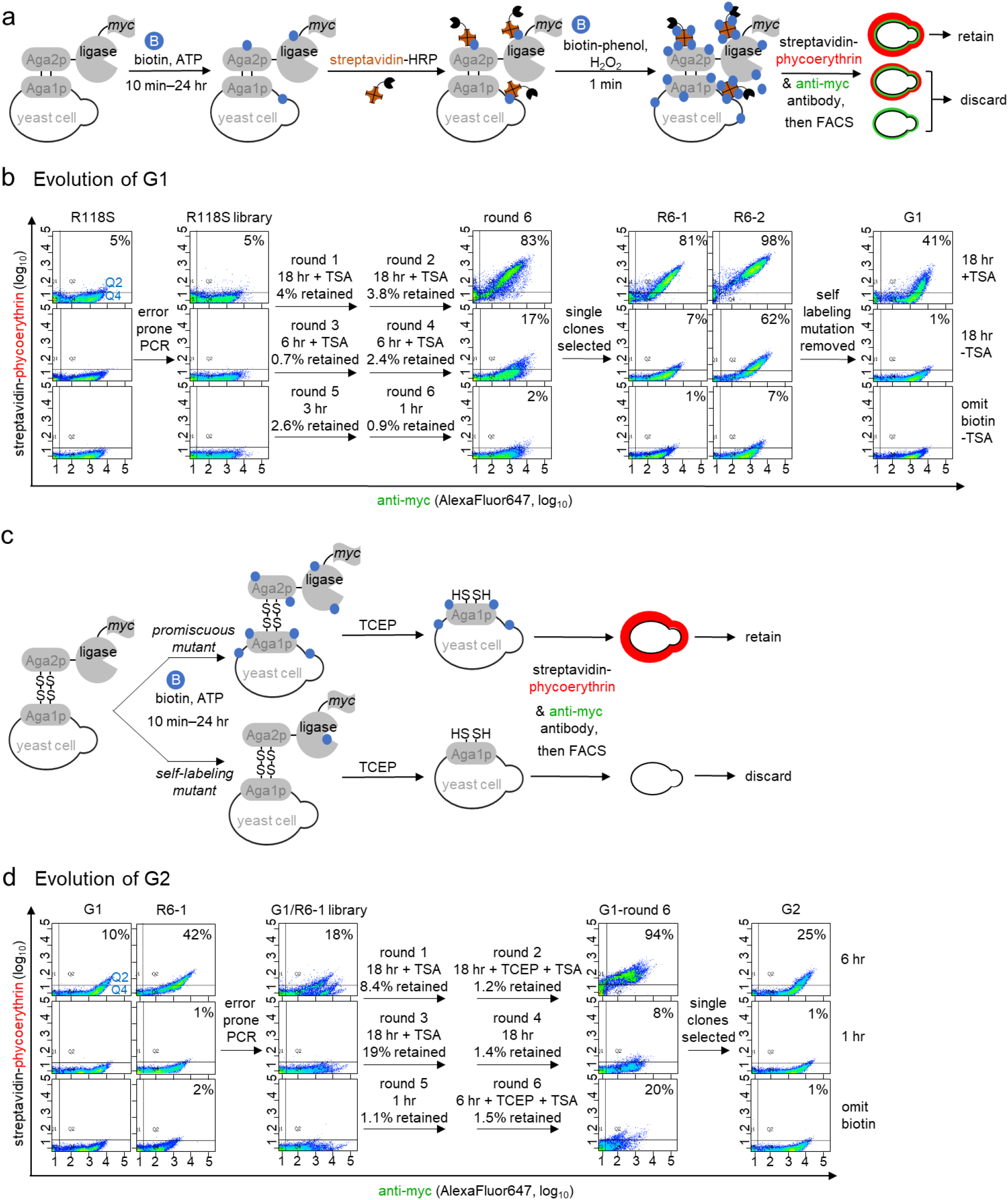
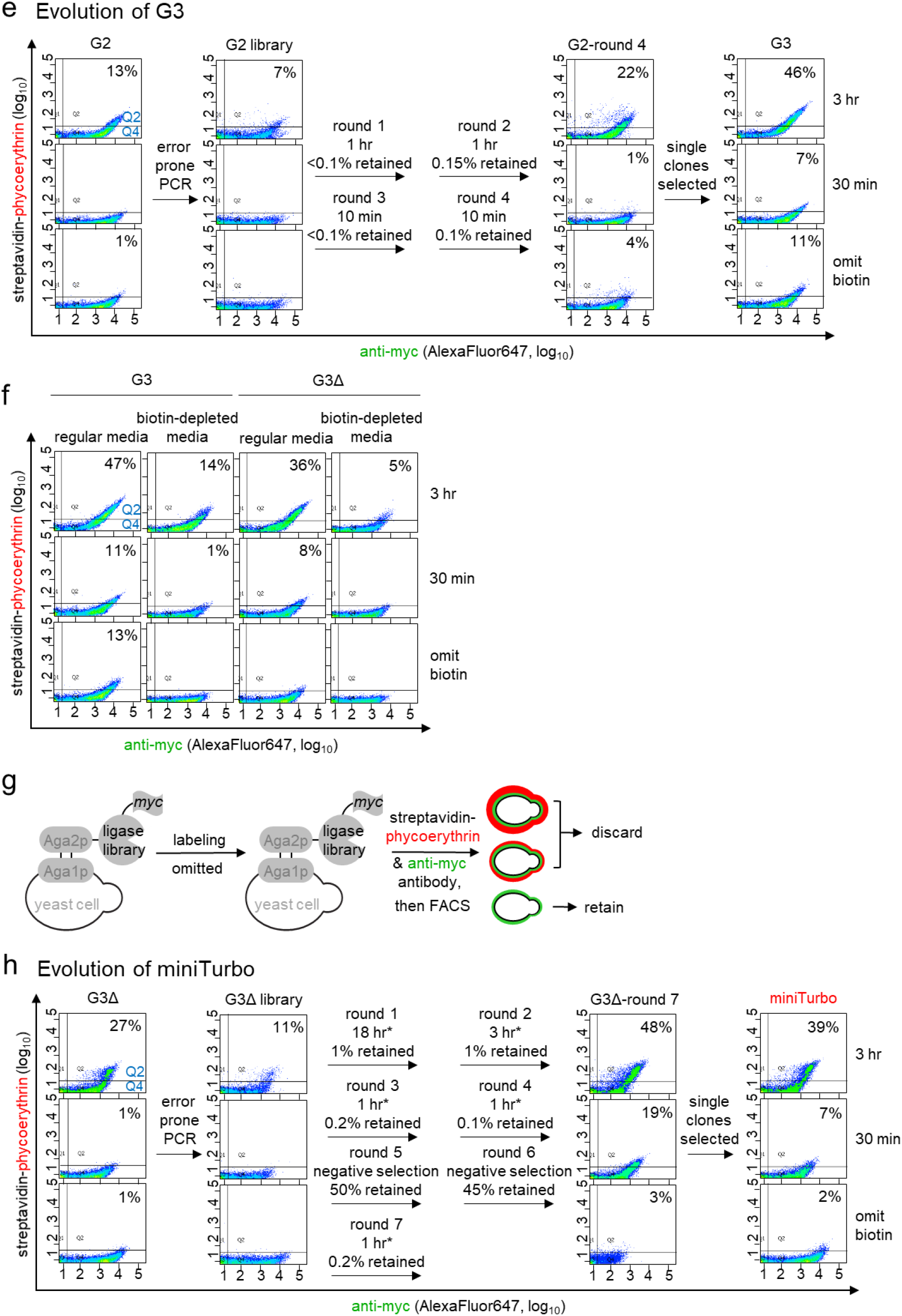
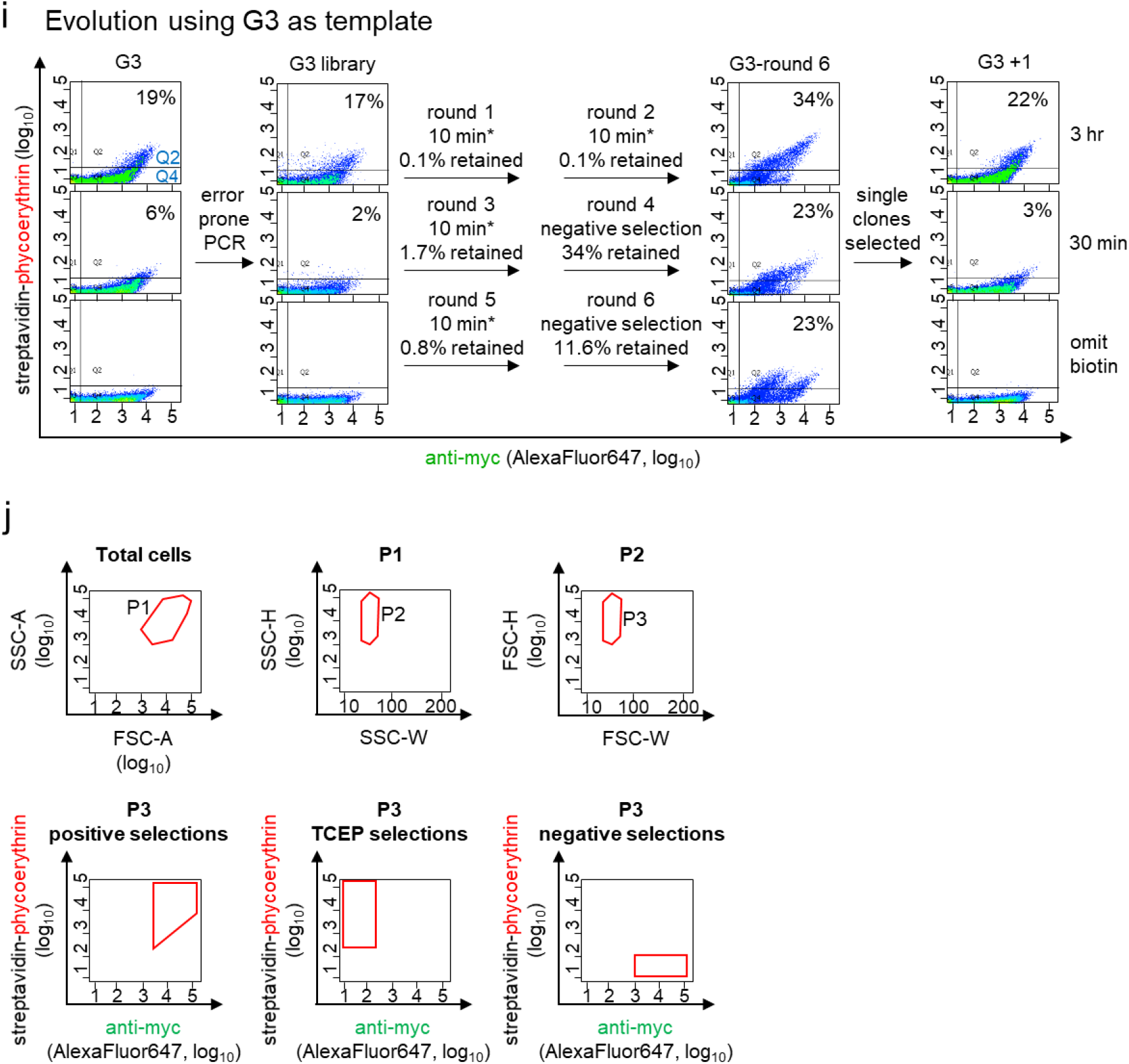
Evolution of TurboID and miniTurbo on yeast. (**a**) Employing TSA to amplify biotinylation signal on the yeast surface. Yeast are labeled with 50 μM biotin and 1 mM ATP for 10 – 24 hr. Prior to staining with fluorophores, yeast are stained with streptavidin-HRP. HRP labeling is then carried out on the yeast surface by addition of biotin phenol and hydrogen peroxide to biotinylate many more sites on the yeast surface^2^. Yeast are then stained with s to visualize biotinylation and anti-myc to quantify ligase expression. Yeast are then sorted using FACS sorting to enrich cells displaying a high ratio of streptavidin to myc staining. (**b**) Evolution of G1 clone from starting template, BirA-R118S. Selection conditions used in rounds 1-6 are shown. All clones and pools were analyzed here in parallel, under three different conditions: 18 hours of treatment with 50 μM biotin and 1 mM ATP, followed by TSA signal amplification as in (a) (top row); same but without TSA signal amplification (middle row); or no treatment and no amplification (bottom row). Cells were stained with streptavidin-phycoerythrin to detect biotinylation sites (y axis) and anti-myc antibody to quantify ligase expression level (x axis). On the top right of each FACS plot is the percentage of cells in quadrant Q2 divided by the sum of cells in Q2 + Q4; where absent, this value is <0.5%. After round 6, two clones were selected, R61 and R62. A self-labeling mutation was removed from R6-2 to give clone G1. All plots display 10,000 cells. This experiment was performed once, but similar results under the same conditions have been replicated at least once for each individual sample in separate experiments. (**c**) Employing TCEP treatment to de-enrich self-labeling mutants. Yeast are labeled with 50 μM biotin and 1 mM ATP for 10 – 24 hr. Prior to staining with fluorophores, yeast are treated with TCEP to reduce the disulfide bonds through which the Aga2p-ligase fusion is attached to the yeast surface, allowing the ligase to be removed from yeast and washed away. TSA as in (a) can be employed after TCEP treatment to amplify remaining biotinylation signal on yeast surface. Yeast are then stained with streptavidin-phycoerythrin to visualize biotinylation and anti-myc to quantify ligase expression. Yeast are then sorted using FACS sorting to enrich cells with streptavidin signal, which represent mutants with promiscuous labeling capabilities. (**d**) Evolution of clone G2 from clone G1. Presentation is the same as in panel (b). Selection conditions used in rounds 1-6 shown, with rounds 2 and 6 employing TCEP to de-enrich self-labeling mutants as in (c). The three analysis conditions used here are 6 hr, 3 hr, or 0 hrs of 50 μM biotin and 1 mM ATP. All plots display 10,000 cells. This experiment was performed once, but similar results under the same conditions have been replicated at least once for each individual sample in separate experiments. (**e**) Evolution of G3 from G2. Same presentation as in (b). Selection conditions used in rounds 1-4 shown. The three analysis conditions used here are 3 hr, 30 min, or 0 min of 50 μM biotin and 1 mM ATP. This experiment was performed once, but similar results under the same conditions have been replicated at least once for each individual sample in separate experiments. (**f**) FACS plots showing that G3 gives biotinylation in the absence of exogenous biotin. When G3 yeast are cultured in biotin-depleted media, this signal is no longer detected (second column). Deletion of the N-terminal domain to give G3Δ reduces biotin affinity and consequently biotinylation activity prior to exogenous biotin addition (third and fourth columns). All plots display 10,000 cells. This experiment was performed once, but similar results under the same conditions have been replicated at least once for each individual sample in separate experiments. (**g**) Employment of negative selections to de-enrich mutants that carry out biotinylation in the absence of exogenous biotin. After ligase expression is induced, labeling with exogenous biotin and ATP is omitted. Yeast are then stained with streptavidin-phycoerythrin to visualize biotinylation and anti-myc to quantify ligase expression. Yeast are then sorted using FACS sorting to enrich cells with high anti-myc signal but without streptavidin signal. (**h**) Evolution of miniTurbo from G3Δ. Same presentation as in (b). Seven rounds of selection were performed, with rounds 5 and 6 being negative selections to remove clones able to carry about biotinylation in the absence of exogenous biotin addition as in (g). Asterisks denote selections performed in biotin-depleted media. The three analysis conditions used here are 3 hrs, 30 min, or 0 hrs of 50 μM biotin and 1 mM ATP, all in biotin-depleted media. All plots display 10,000 cells. This experiment was performed once, but similar results under the same conditions have been replicated at least once for each individual sample in separate experiments. (**i**) Evolution using G3 as starting template. Same presentation as in (b). Six rounds of selection were performed, with rounds 4 and 6 being negative selections as in (g) to remove clones that are able to carry out biotinylation in the absence of exogenous biotin addition. Asterisks denote selections performed in biotin-depleted media. The three analysis conditions used here are 3 hrs, 30 min, or 0 hrs of 50 μM biotin and 1 mM ATP, all in biotin-depleted media. All plots display 10,000 cells. This experiment was performed once, but similar results under the same conditions have been replicated at least once for each individual sample in separate experiments (except G3 + 1 which was only performed once). (**j**) Examples of various gates drawn for FACS sorting. Top 3 plots indicate gates used to analyze single-cell yeast populations, bottom 3 plots indicate gates used to retain yeast for positive selections (left), positive selections using TCEP (middle), and negative selections (right). X and y-axes indicated for each plot. Gates are drawn in red with the relevant resulting population written next to it. SSC-A is side-scatter area, FSC-A is forward-scatter area, SSC-H is sidescatter height, SSC-W is side-scatter width, FSC-H is forward-scatter height, FSC-W is forward-scatter width.

**Supplementary Figure 3.**
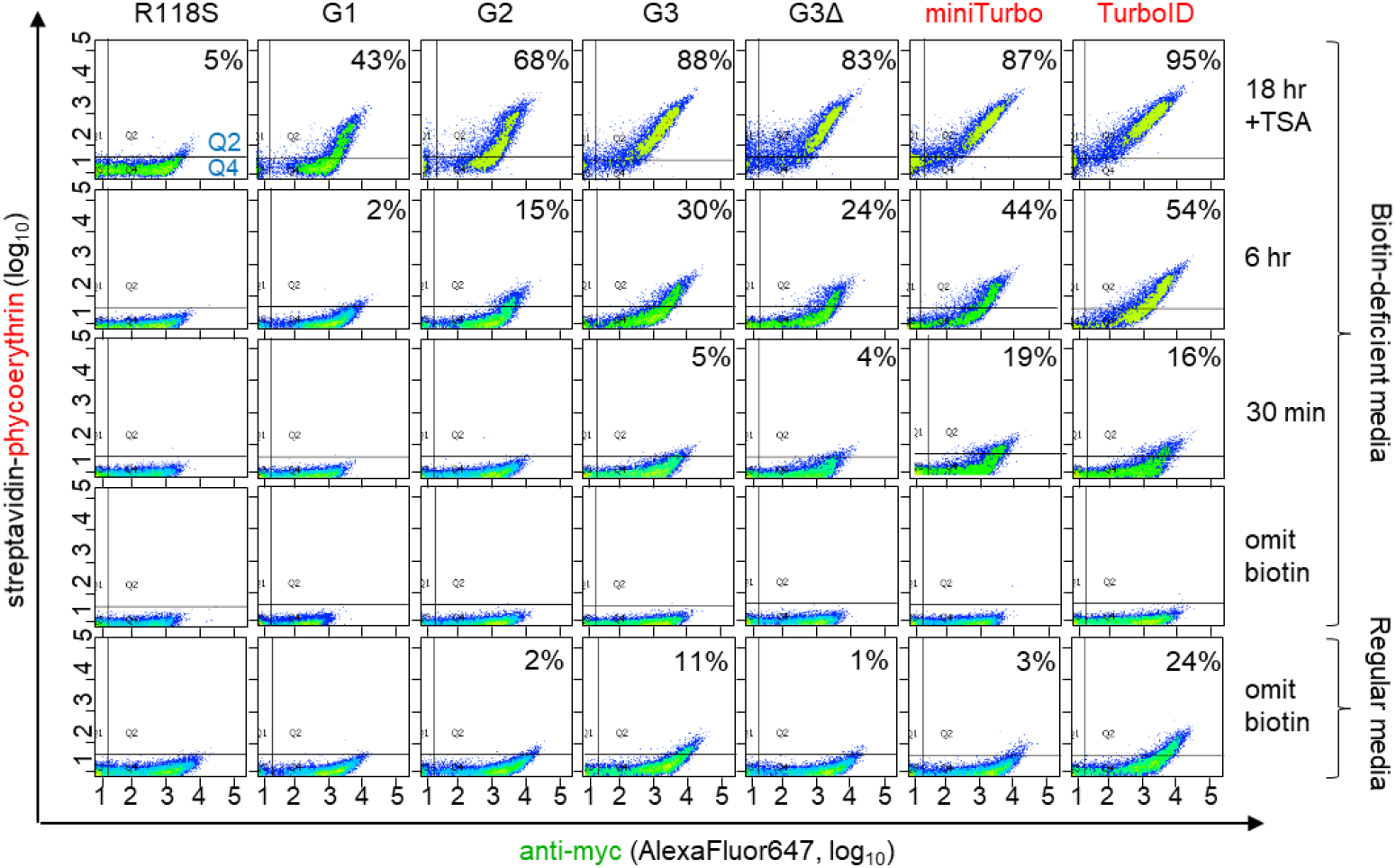
FACS plots summarizing progress of evolution. Same as Figure 1e, but with more time points. In addition to 6 hours of labeling with 50 μM biotin, the results of 18 hr and 30 min labeling are shown. The first four rows were carried out in biotin-depleted media, while the last row was in regular yeast media. All plots display 10,000 cells. This experiment was repeated one time (except for G3Δ and the “biotin omitted” conditions in “biotin-deficient media” which were performed only once under these conditions).

**Supplementary Figure 4.**
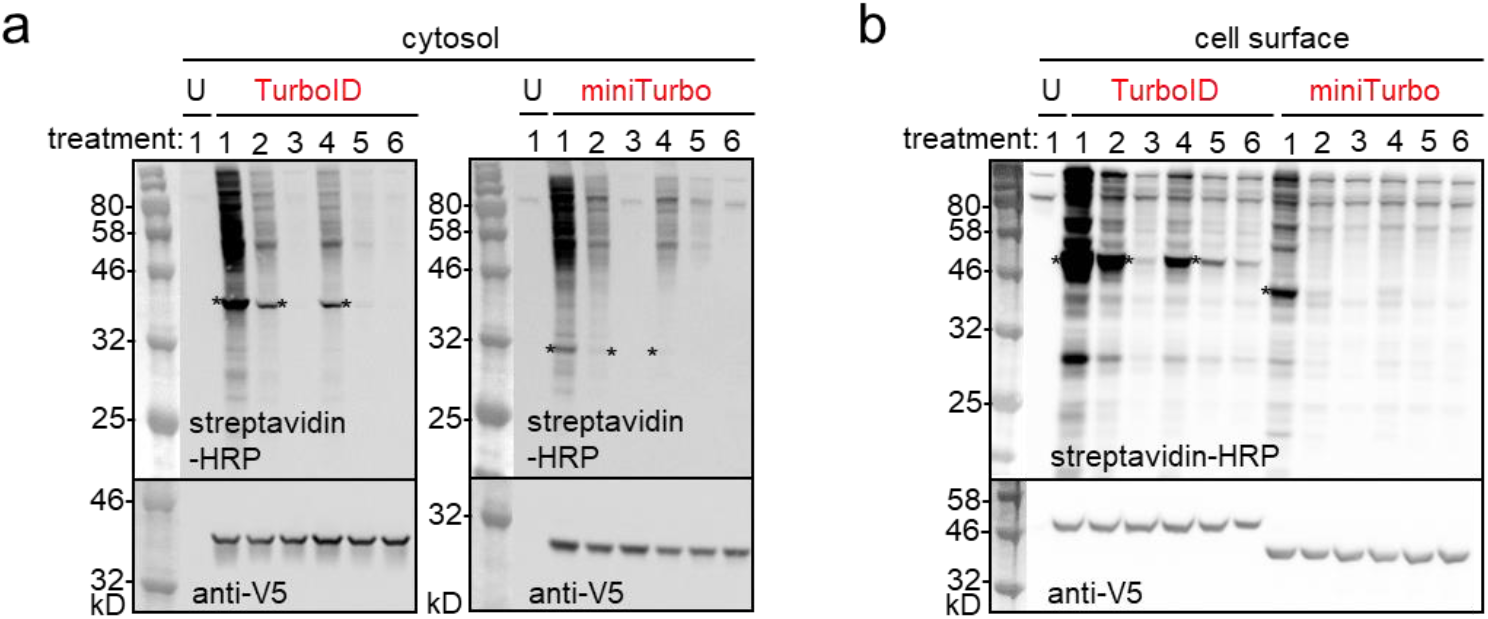
Halting promiscuous labeling by TurboID and miniTurbo by lowering temperature to 4°C. The indicated ligases were transiently expressed in HEK 293T cells targeted to (**a**) the cytosol or (**b**) the cell surface. Labeling was carried out under the following conditions as indicated: 1. Incubation with 500 μM biotin for 70 min at 37°C. 2. Incubation with 500 μM biotin for 10 min at 37°C. 3. No biotin incubation. 4. Incubation with 500 μM biotin for 10 min at 37°C, then moved to 4°C for 60 more min. 5. Incubation with 500 μM biotin for 60 min at 4°C. 6. Incubation with 500 μM biotin for 10 min at 4°C. For (b), samples were also incubated with 0.5 mM ATP and 1.25 mM magnesium acetate. After labeling, whole cell lysates were analyzed by streptavidin blotting. Ligase expression detected by antiV-5 blotting. U, untransfected. Asterisks indicate ligase self-biotinylation. This experiment was performed once, but similar results were replicated for BirAG3.

**Supplementary Figure 5.**
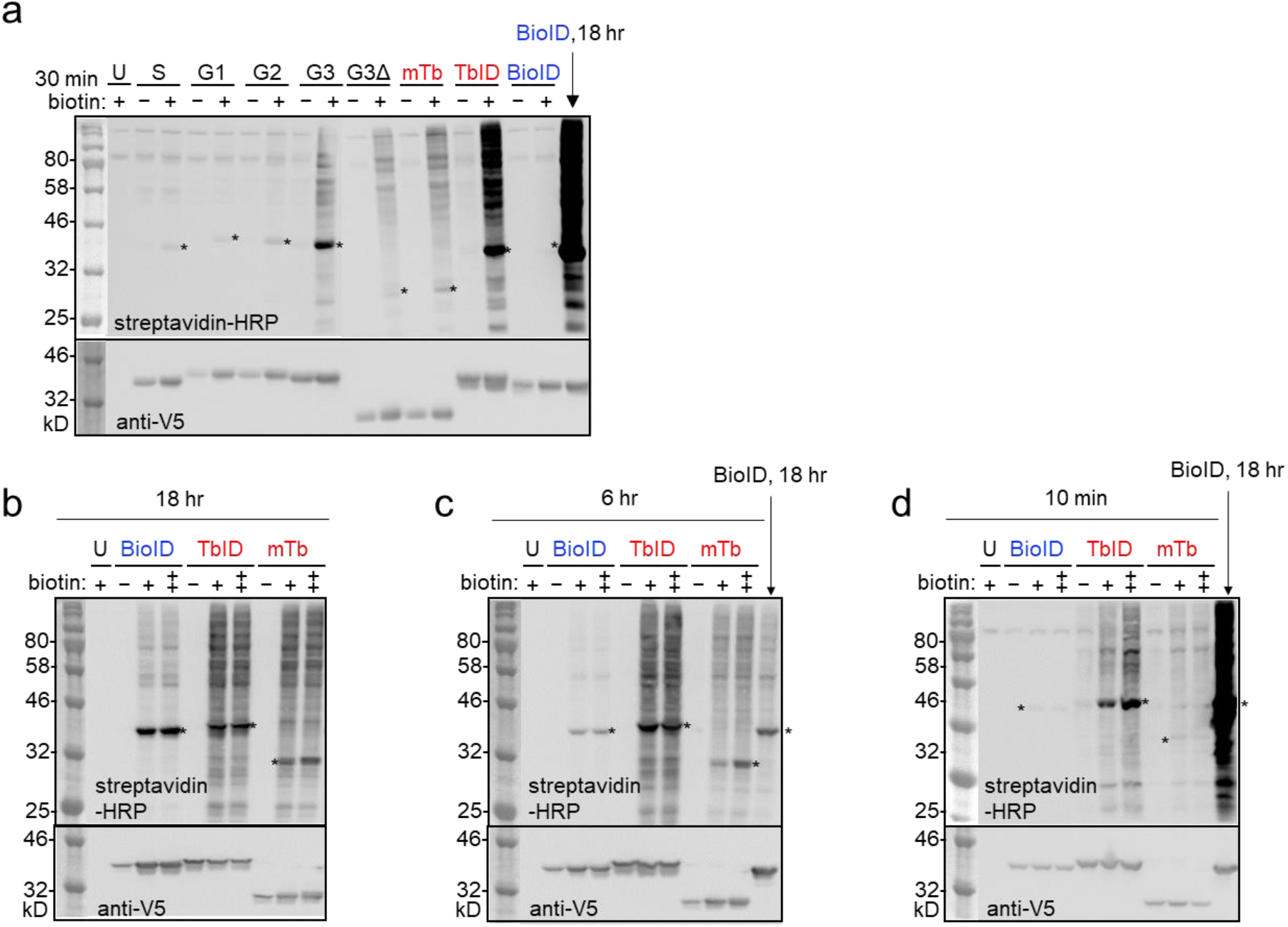
Comparison of ligase activities in HEK cytosol. (**a**) The indicated ligases were transiently expressed in the cytosol of HEK 239T cells. 50 μM exogenous biotin was added for 30 minutes, then whole cell lysates were analyzed by streptavidin blotting. Ligase expression detected by anti-V5 blotting. U, untransfected. S, BirA-R118S. Asterisks indicate ligase self-biotinylation. BioID labeling for 18 hours shown in the last lane for comparison. This experiment was performed once. (**b**)-(**d**) Same as (a) but with different labeling times, as indicated. In the “++” lanes, 500 μM exogenous biotin was added to cells. These experiments were performed once, but has been repeated for BioID under 6 hr labeling conditions twice; and BioID under 18 hr labeling conditions ([b]) and TurboID and miniTurbo under 10 min labeling conditions ([d]) have been repeated more than three times.

**Supplementary Figure 6.**
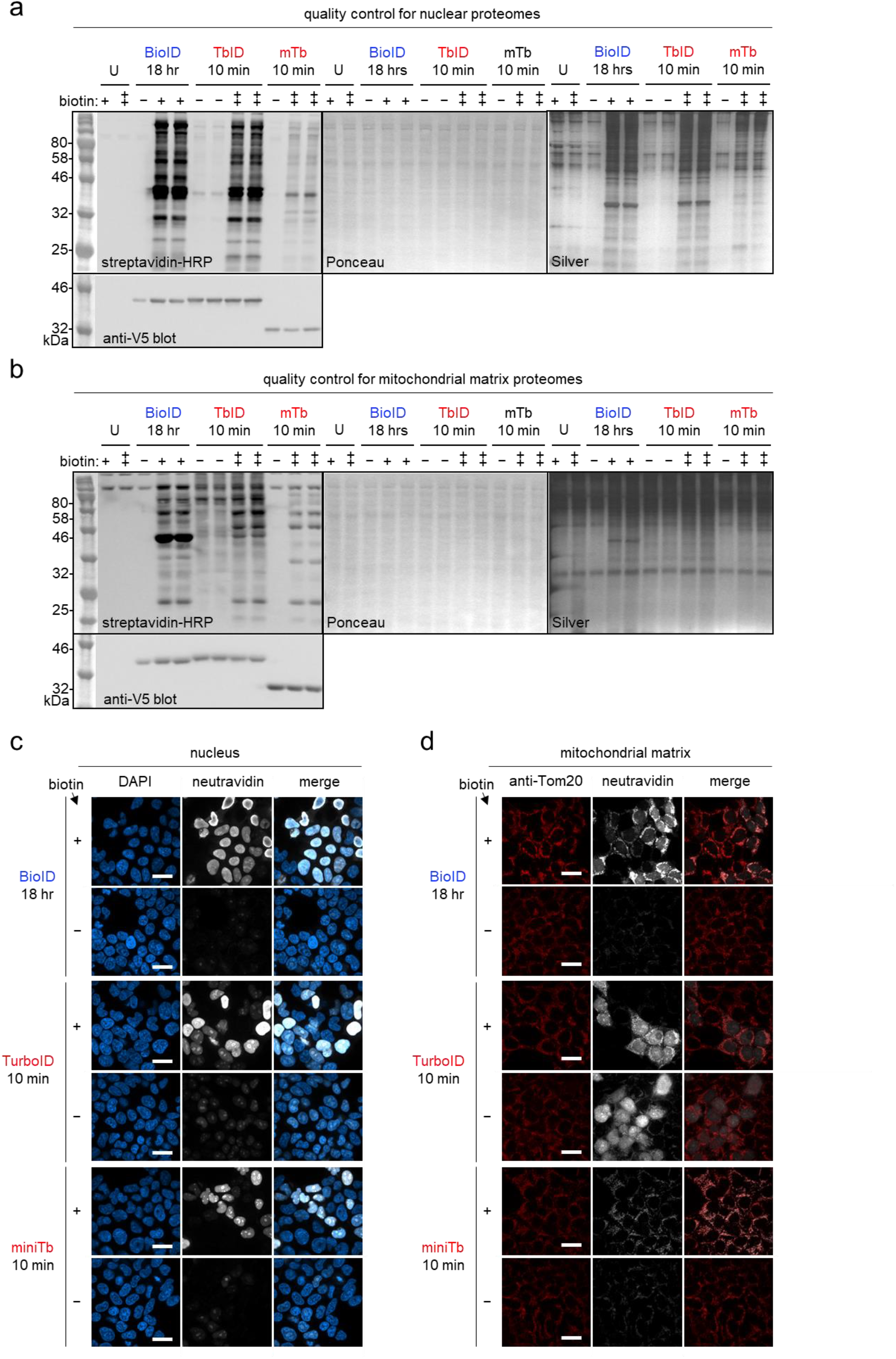
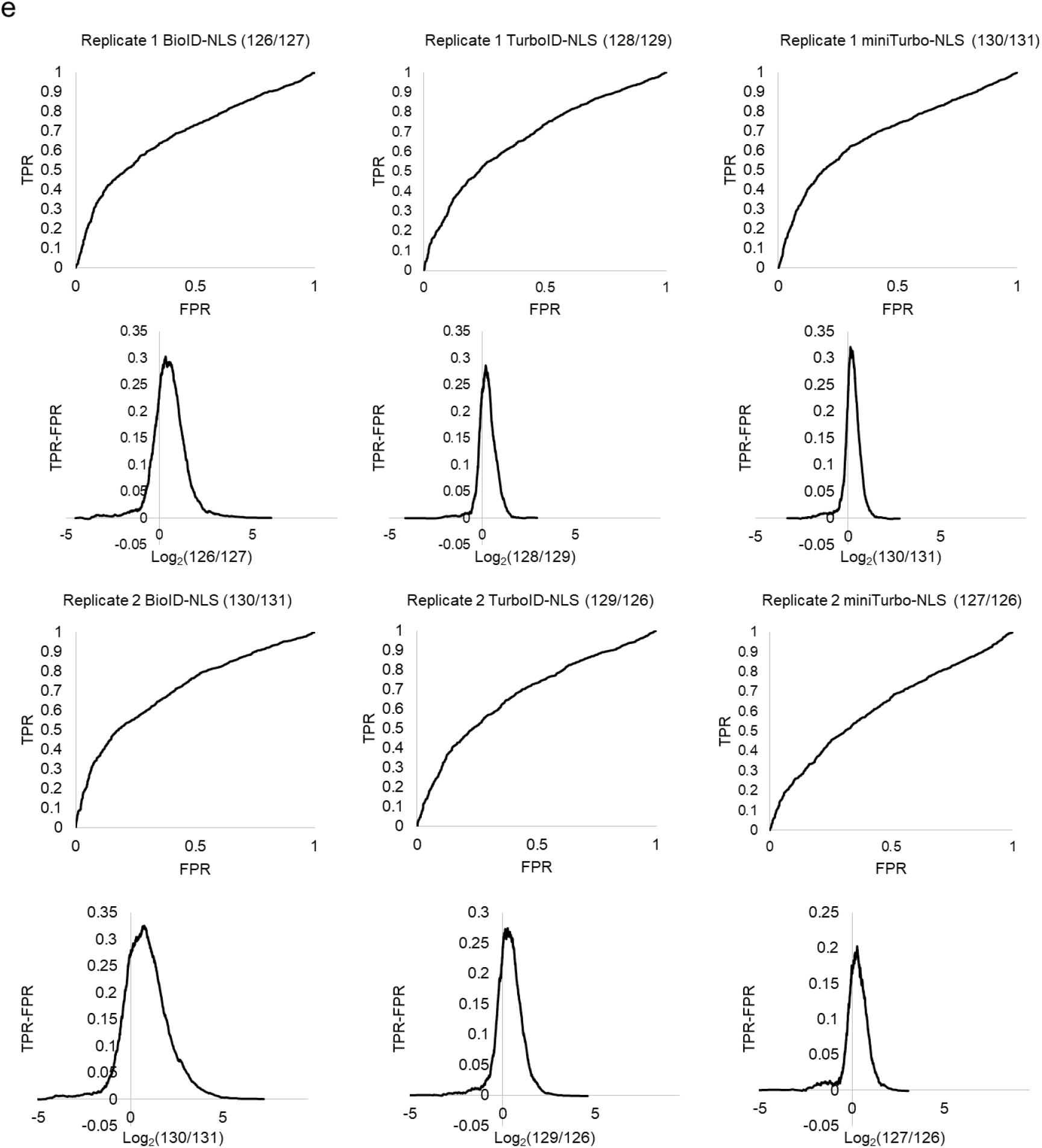
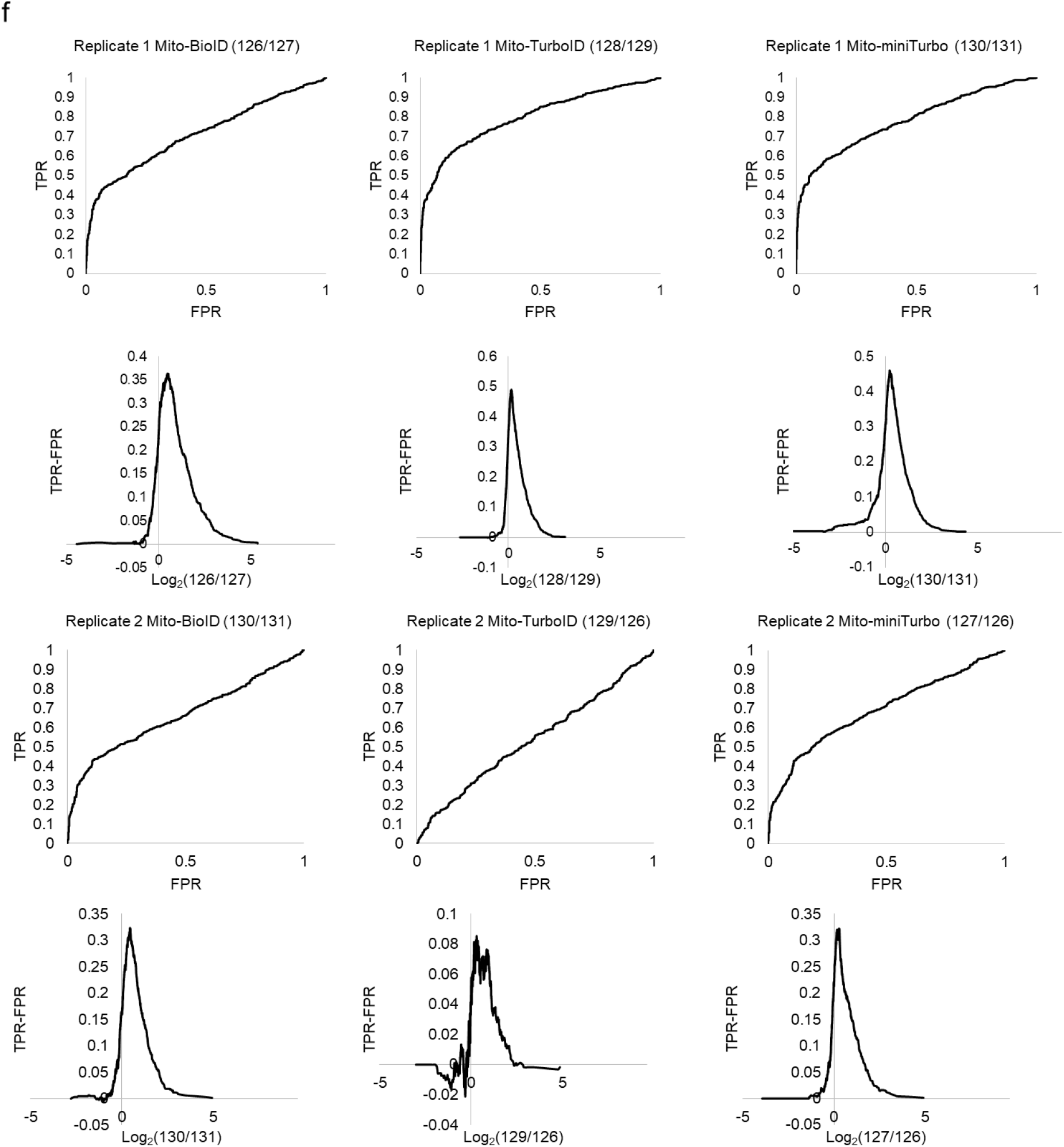
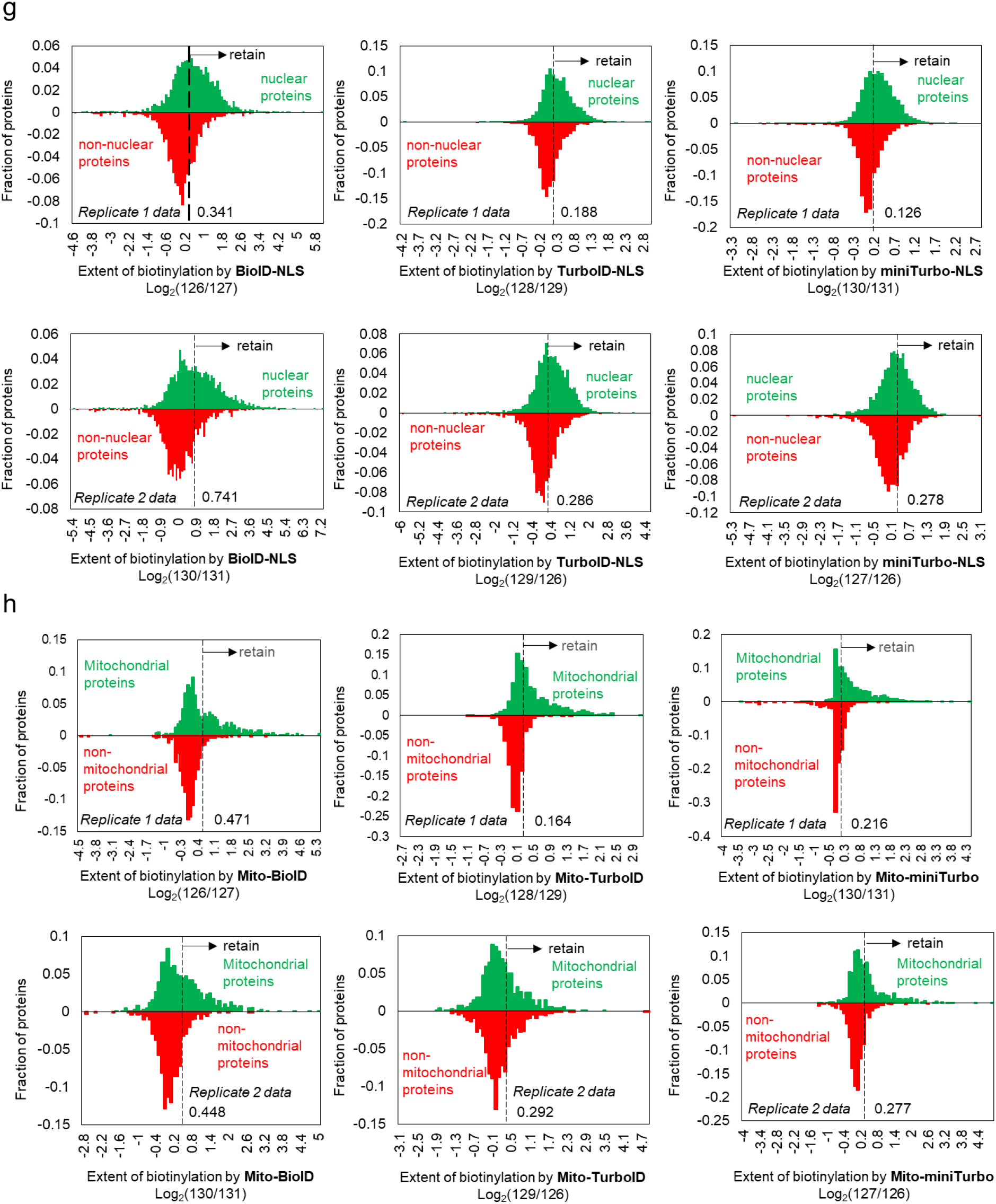
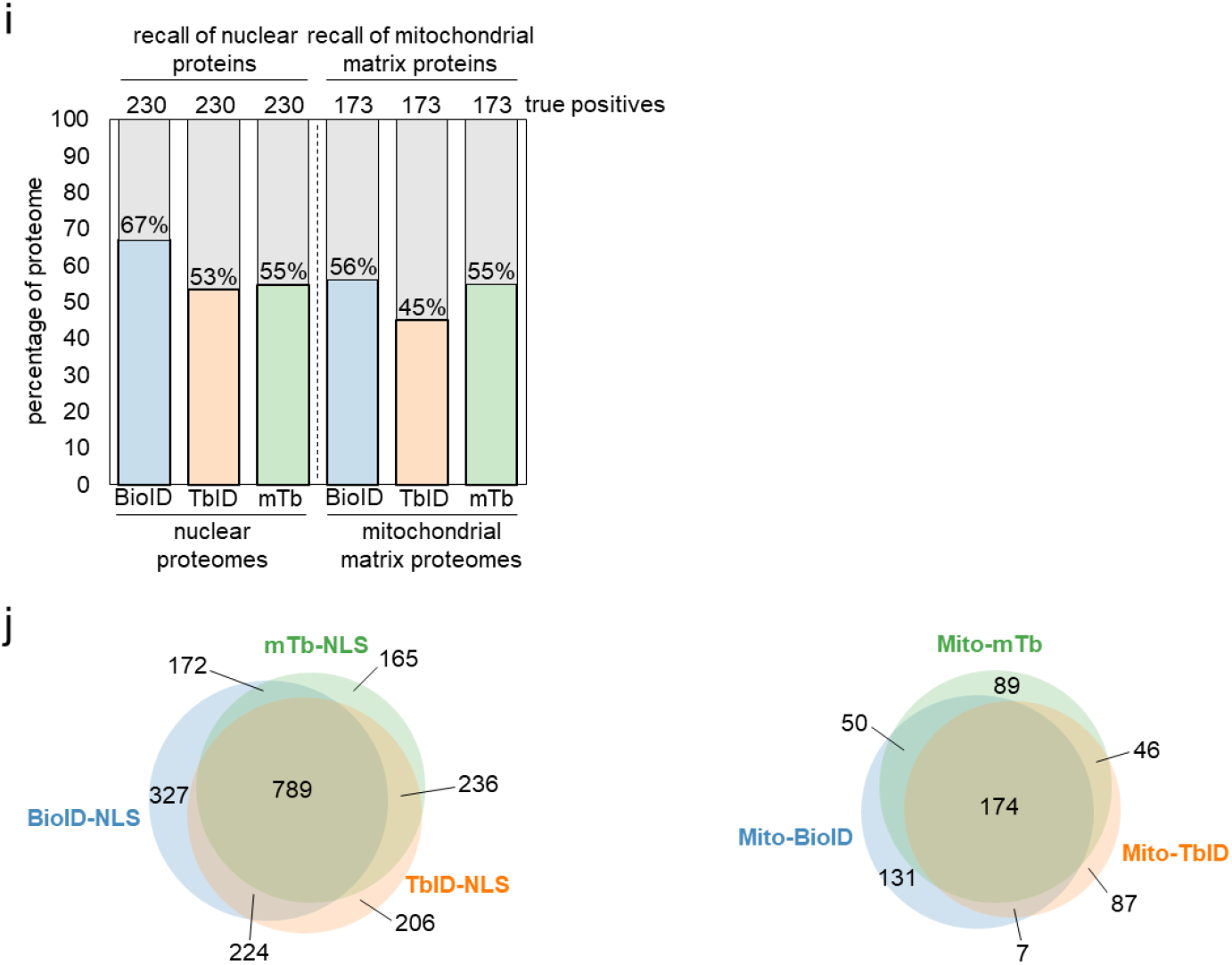
Mass spectrometry-based proteomic experiment using TurboID, miniTurbo, and BioID. **(a, b)** Quality control gel analysis of (a) nuclear and (b) mitochondrial matrix proteomic samples. 2.5% of whole cell lysate was used for analysis by streptavidin blotting and ligase expression detection by anti-V5 staining (left). Ponceau staining (middle) shows equal loading of samples. Right: 5% of streptavidin beads were boiled in SDS buffer to elute biotinylated proteins. Eluted proteins were separated on SDS-PAGE and detected by silver stain (right). The proteomics experiment was performed once with two replicates for each construct, but small scale pulldown experiments were repeated once for both nuclear and mitochondrial matrix samples. (**c, d**) Characterization of biotin-labeled cells by fluorescence microscopy. HEK prepared and labeled as in Figure 2d were fixed and stained with neutravidin-AlexFluor647 to visualize biotinylated proteins. DAPI was also used to stain nuclei in (c), and anti-Tom20 antibody to stain mitochondria in (d). Scale bars, 20 μm. In the BioID and TurboID mitochondrial samples, we also observe some neutravidin staining in the cytosol and nucleus, which we believe results from (1) some mis-targeting of subpopulations of each ligase, perhaps due to promiscuous biotinylation of N-terminal mitochondrial targeting sequences (which contains lysine), which may impair mitochondrial import, and (2) ligases being more active in the cytosol and nucleus than in the mitochondrial matrix. (**e**) Determination of optimal TMT ratio cutoffs for both replicates of nuclear proteomes. For every possible TMT ratio cutoff, the true positive rate (TPR) was plotted against the false positive rate (FPR) in a receiver operating characteristic (ROC) curve (top). TPR is defined as the fraction of detected true positive proteins above the cutoff. FPR is defined as the fraction of detected false positive proteins above the cutoff. The bottom graph plots the difference between the TPR and FPR at every TMT ratio cutoff. Cutoff is made at the TMT ratio at which TPR-FPR is maximal, and it is depicted in the histogram in (g) as a dashed line. (**f**) Same as (e), but for mitochondrial proteomes. (**g**) Histograms that illustrate how cutoffs were determined to identify proteins biotinylated by the indicated ligase. True positives (i.e., nuclear annotated proteins) are plotted in the green histogram, and potential false positives are plotted in red histograms. Receiver-operating characteristic analysis from (e) was applied to determine the TMT ratio cutoff value that maximized true positives while minimizing false positives. (**h**) Same as (g), but for mitochondrial proteomes. (**i**) Coverage analysis for each proteomic dataset. Graph indicates the fraction of true positive proteins recalled in each proteome. True positive proteins used for this analysis were curated from literature. (**j**) Overlap of BioID (blue), TurboID (orange) and miniTurbo (green) derived proteomes for the nucleus (left) and mitochondrial matrix (right). Proteins that were only labeled by BioID or proteins that were only labeled by TurboID or miniTurbo were less specific for the compartment being mapped. For example, for the mitochondrial matrix proteomes, only 33% of BioID-only detected proteins and 35% of TurboID/miniTurbo-only detected proteins had previous mitochondrial annotation, whereas 93% of proteins detected by all three ligases had previous mitochondrial annotation; and for the nucleus, 71% of BioID-only detected proteins and 62% of TurboID/miniTurbo-only detected proteins had previous nuclear annotation, whereas 84% of proteins detected by all three ligases had previous nuclear annotation.

**Supplementary Figure 7.**
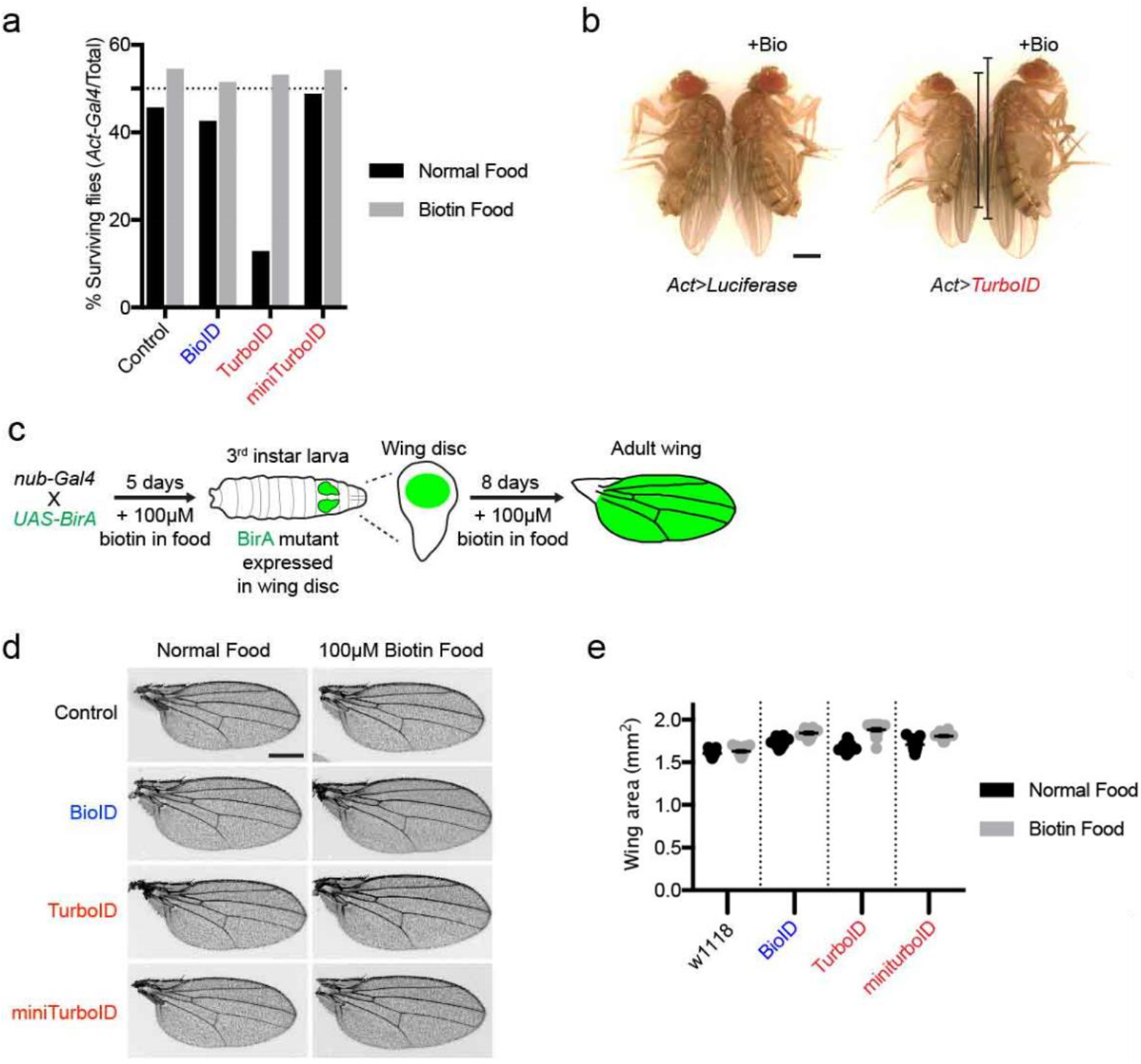
Viability and morphology assays in *D. melanogaster* expressing promiscuous biotin ligase variants. (**a**) Viability assay. *Act-Gal4* drives expression of the indicated *UAS-BirA* enzyme in all cells at all developmental time points. In control flies, *Act-Gal4* drives expression of *UAS-luciferase* in all cells at all developmental time points. Graph indicates the percentage of surviving flies that are ubiquitously expressing the indicated BirA enzyme. Dashed line indicates expected frequency of flies expressing BirA (50%). Flies were either raised on control food or biotin food. Biotin food is control food supplemented with 100 μM biotin. Sample size values (n) from left column to right: 512, 286, 586, 524, 466, 513, 563, 459. This experiment was performed twice with similar results, and the counts combined. (**b**) Image of two adult *Act-Gal4, UAS-luciferase* flies and two adult *Act-Gal4, UAS-TurboID* flies from (a) that are grown on control food and control food supplemented with 100 μM biotin (+Bio). Vertical black lines illustrate size difference between flies expressing TurboID that are raised on normal food versus biotin food. Adult flies shown are all female. (**c**) Schematic of tissue specific expression of BirA enzymes in wing imaginal disc. *Nub-Gal4* drives expression of the indicated *UAS-BirA* in the wing disc pouch, which gives rise to the adult wing blade. Flies were provided food supplemented with 100 μM biotin for 13 days after egg deposition, then the wings were dissected from adult flies. (**d**) Images of adult wings as prepared in (c). Control flies are untransfected (*w1118*). Flies were raised on control food, or control food supplemented with 100 μM biotin. Scale bar is 0.5 mm. Adult wings shown are all from females. (**e**) Quantitation of wing area of specimens prepared as described in (c). Control flies are non-transgenic (*w1118*). Biotin food is control food supplemented with 100 μM biotin. Sample size values (n) from left column to right: 17, 14, 17, 15, 19, 18, 19, 18. Error bars in s.e.m. This experiment repeated twice.

